# Litter decomposition rates of biocrust-forming lichens are similar to that of vascular plants and are affected by warming in a semiarid grassland

**DOI:** 10.1101/2020.04.09.019695

**Authors:** Miguel Berdugo, Dinorah O. Mendoza-Aguilar, Ana Rey, Victoria Ochoa, Beatriz Gozalo, Laura García-Huss, Fernando T. Maestre

## Abstract

Despite the high relevance of communities dominated by lichens, mosses and cyanobacteria living on the soil surface (biocrusts) for ecosystem functioning in drylands worldwide, no study to date has investigated the decomposition of biocrust-forming lichen litter *in situ*. Thus, we do not know whether the drivers of its decomposition are similar to those for plant litter (e.g., importance of abiotic degradation through UV radiation), the magnitude of lichen decomposition rates and whether they will be affected by climate change. Here we report results from a litter decomposition experiment carried out with two biocrust-forming lichens (*Diploschistes diacapsis* and *Cladonia convoluta*) in central Spain. We evaluated how lichen decomposition was affected by warming, rainfall exclusion and the combination of both. We also manipulated the incidence of UV radiation using mesh material that blocked 10% or 90% of incoming UV radiation. Our results indicate that lichens decompose as fast as some plants typical of the region (*k*~0.3) and that the chemical composition of their thallus drives litter decomposition rates. Warming increased decomposition rates of both lichen species, and mediated the effects of photodegradation. While UV exposure accelerated the decomposition of *D. diacapsis*, it slowed down that of *C. convoluta*. Our results indicate that biocrust-forming lichens can decompose in the field at a rate similar to that of vascular plants, and that this process will be affected by warming. Our findings further highlight the need of incorporating biocrusts into carbon cycling models to better understand and forecast climate change impacts on terrestrial biogeochemistry.

## 1. Introduction

Arid, semi-arid and dry-subhumid ecosystems (drylands, hereafter) cover over 41% of the terrestrial surface (Cherlet and others 2018), but forecasted increases in aridity will increase their global area by 11-23% by the end of this century (Huang and others 2015). Drylands account for over 25% of the total amount of organic carbon stored in the world’s soils (Safriel and Adeel 2005) and are key to understand the inter-annual variability of the global carbon cycle (Poulter and others 2014). In these areas, an important part of soil carbon is stored in the first soil centimetres (Ciais and others 2011; Thomas 2012). Thus, soil carbon dynamics in drylands is largely affected by those organisms and communities living on the soil surface, such as biocrusts (communities constituted by cyanobacteria, algae, fungi, lichens, bryophytes and other microorganisms living in an intimate association with soil particles; Belnap & Lange 2013).

Biocrusts are known to affect multiple processes of the carbon cycle. They fix atmospheric CO_2_ and increase the amount of carbon stored in soils (Darrouzet-Nardi and others 2015; Delgado-Baquerizo and others 2015a), modulate carbon losses to the atmosphere via soil respiration (Castillo-Monroy and others 2011; Thomas 2012; Escolar and others 2015), and affect the activity of soil enzymes such as β-glucosidase (Bowker and others 2011; Miralles and others 2013). Despite their global extent, it is estimated that biocrusts cover over 12% of terrestrial surface (Rodriguez-Caballero and others 2018), and impacts on the carbon cycle, their role on processes such as litter decomposition is virtually unknown in drylands.

Litter decomposition is a key process in the global carbon cycle, as more than 50% of net primary production returns to soils via the decomposition of plant litter in terrestrial ecosystems (Wardle and others 2004). Multiple studies have shown that the decomposition of lichen and moss tissues is an important input of carbon and nutrients into the soil in boreal/sub-boreal (Wetmore 1982; Lang and others 2009; Campbell and others 2010) and temperate (McCune and Daly 1994; Caldiz and others 2007; Li and others 2014) ecosystems. Indeed, the rates at which bryophytes and lichens decompose in these systems can be as high as that of some vascular plants (Lang and others 2009). Studies conducted in artic areas have found important differences in the decomposition rates of several lichen species (Lang and others 2009), which are generally attributed to the thallus nutrient content (fast decomposition for species with high N and low C to N ratio) and growing form (faster for fruticose cf. foliose). However, no previous study has, to our knowledge, evaluated in the field the decomposition of biocrust-forming lichens in drylands. As such, the rate at which their tissues decompose, and the contribution of this process to carbon cycling in these areas, is unknown.

In drylands, the rate at which plant litter is decomposed depends on the interaction between litter chemical composition and multiple biotic and abiotic factors, including soil nutrients, soil fauna and microorganisms and climatic conditions such as precipitation and UV radiation (McCulley and others 2005; Throop and Archer 2009; Brandt and others 2010; King and others 2012; García-Palacios and others 2013; Delgado-Baquerizo and others 2015b; Almagro and others 2017). In particular, a growing number of studies have shown that solar ultraviolet (UV) radiation (280-400 nm) can be an important driver of leaf litter decomposition in dryland ecosystems by directly breaking down organic matter components releasing CO_2_, a process called photodegradation (Brandt and others 2007; Day and others 2007; Gallo and others 2009; Rutledge and others 2010). Although results differ among litter types, site characteristics (solar irradiance dose, temperature, moisture, etc.) and experimental conditions (field vs. laboratory) (Austin and Vivanco 2006; Gallo and others 2006; Brandt and others 2010; Almagro and others 2017), a recent meta-analysis concluded that solar radiation speeds up decomposition by 32% (King and others 2012). Because of the lack of studies on this topic conducted in drylands, we do not know whether photodegradation may also contribute to the decomposition of biocrust-forming lichens in these environments.

The decomposition of biocrust constituents may be more important than it seems, particularly in the face of climate change. Multiple experimental studies have shown that increases in temperature and/or alterations in precipitation such as those forecasted by climatic models reduce the photosynthetic performance, growth and cover of biocrust constituents such as mosses and lichens (Maphangwa and others 2012; Reed and others 2012; Ladrón de Guevara and others 2014; Ferrenberg and others 2015; Maestre and others 2015). There is also experimental evidence that warming-induced changes in biocrust communities may act synergistically with other climate change-induced effects, such as the increase in soil CO_2_ efflux and alterations in bacterial and fungal communities, altering soil carbon cycling and reducing soil carbon storage in drylands (Maestre and others 2013). For instance, increases in surface soil carbon with warming despite the reductions in the photosynthetic activity and cover of biocrust-forming lichens have been reported in climate change experiments conducted in central and south-east Spain (Maestre and others 2013; Ladrón de Guevara and others 2014; García-Palacios and others 2018). Maestre and others (2013) observed that increases in recalcitrant carbon compounds in the soil, which are abundant in the lichen thalli, were positively correlated with biocrust cover losses, suggesting that the decomposition of dead lichens due to warming could explain the observed increase in soil carbon contents. These findings highlight the need to improve our knowledge of the decomposition rate of biocrust components such as lichens, and indicate that understanding litter decomposition processes within biocrusts is essential for improving our understanding of carbon cycling in drylands under climate change. Such knowledge would constitute a first step towards developing more realistic and accurate carbon dynamic models incorporating biocrusts, which right now are largely absent from the suite of models of carbon cycling being currently used (Resplandy and others 2018).

Here we evaluate, for the first time, how simulated climate change (3°C warming and 35% rainfall reduction) and UV radiation affect the decomposition of biocrust-forming lichen thalli under field conditions in a semiarid grassland from central Spain. We compared the effects of UV radiation and climate manipulation treatments on the decomposition rate of two dominant species in our study area, *Cladonia convoluta* (Lam.) Anders and *Diploschistes diacapsis* (Ach.) Lumbsch. Using a manipulative experiment conducted over 2.5 years, we aimed to: i) quantify the litter decomposition rates of *Cladonia* and *Diploschistes*, ii) investigate how litter quality affects these rates, iii) assess the effects of simulated climate change (warming and rainfall reduction) on the decomposition rate and chemistry of lichen litter, and iv) investigate whether UV radiation (photodegradation) accelerates its decomposition. We tested the following hypotheses: i) the two species studied will exhibit different decomposition rates due to differences in their tissue chemistry (Lang and others 2009), ii) litter lichen decomposition rate will be reduced by warming and reduced precipitation due to lower water availability (Almagro and others 2015), and iii) the relative contribution of photodegradation will increase in response to climate change, as previously observed with plant litter in our study area (Almagro and others 2017).

## 2. Material and Methods

### 2.1. Study area

The study was carried out in the Aranjuez Experimental Station, located in central Spain (Figure S1) (40°02’ N–3°32’ W; 590 m a.s.l). The climate is Mediterranean semi-arid, with mean annual temperature and rainfall values of 15°C and 349 mm, respectively. Precipitation events predominantly occur in autumn-winter and spring. The soil is classified as Gypsiric Leptosol (IUSS Working Group WRB 2015). The cover of perennial plants is lower than 40%, and is dominated by the perennial herbaceous species *Macrochloa tenacissima*. To a lesser extent, there are also isolated individuals of shrubs such as *Helianthemum squamatum, Gypsophila struthium* and *Retama sphaerocarpa*. Open areas between plants contain a well-developed biocrust community dominated by lichens such as *Diploschistes diacapsis*, *Squamarina lentigera*, *Fulgensia subbracteata* and *Buellia zoharyi*. Mosses such as *Pleurochaete squarrosa* and *Didymodon acutus* are also present (mostly under the canopy of *Macrochloa*, where *Cladonia convoluta* can also be found) at the study site, as well as cyanobacteria of the genera *Microcoleus*, *Schizothrix*, *Tolypothrix*, *Scytonema* and *Nostoc* (Cano-Díaz and others 2018). See Maestre et al. (2013) for a complete species checklist of biocrust-forming moss and lichens found at our study site.

### 2.2. Lichen litter collection

For this study, the selected species were *Diploschistes diacapsis*, a crustose lichen, and *Cladonia convoluta*, a fruticose lichen (Figure S2). Both species have preference for gypsum soils and are semi-vagrants or vagrants lichens. They are also common in the study area, and due to their loose attachment to the soil can be collected without disturbing the soil surface nor affecting other biocrust-forming species. Thalli from *D. diacapsis* and *C. convoluta* were collected in June 2013 at the study area and were transported to the laboratory, where they were cleaned by removing dirt and other cryptogam species. Once cleaned, we killed living tissues by keeping them frozen at −80° C for a week; after that, the thalli were submerged in liquid nitrogen for nearly 5 seconds (see Lang and others, 2009 for a similar approach). After these treatments, the thalli were oven-dried at 70° C for 24 hours.

To study the decomposition of lichen thalli in the field, we used the litter bag method (Lang and others 2009; García-Palacios and others 2013). Samples of lichen-litter (~1.6 g per litterbag) were weighted and placed in 5 x 5 cm litterbags with a mesh size of 1 mm^2^ (Figure S2 C). We used two types of mesh material to build the litterbags (Light treatment): a UV-block screen material made of fibreglass, which blocked ~90% of incoming UV-radiation, and a UV-pass screen material made of high-density polyethylene, which blocked only ~10% of incoming UV-radiation (Dirks and others 2010; Almagro and others 2015).

### 2.3. Experimental design

To evaluate the impacts of climate change on the decomposition rates of lichen litter, and the interaction effects with photodegradation, a factorial experiment was set-up with four climatic treatments: *warming* (W, a ~2.5°C annual temperature increase), *rainfall reduction* (RE, a ~35% reduction in annual rainfall), the combination of both *warming and rainfall reduction* (WRE) and a *control* (C) treatment with no manipulation. The experimental design consisted of five blocks distributed among the different climate manipulation treatments in a split-plot design (20 whole plots and 80 subplots in total, Figure S3). The whole-plot factor was the different climatic manipulations used (C, W, RE and WRE). The subplot factor was a factorial combination of Light treatment (UV-block, UV-pass) and litter species (*D. diacapsis* and *C. convoluta*), which were randomly assigned within the plots. The experiment was set up in early June 2013.

### 2.4. Climate manipulation treatments

To increase temperature in the warming treatment, we used open top chambers (OTCs) with hexagonal design and sloping sides of 65 × 42 × 52 cm. These chambers were built with methacrylate sheets, which ensure high transmittance in the visible spectrum (92%, according to the manufacturer; Decorplax S.L., Humanes, Spain) and a very low emission in the infrared wavelength. Chambers were suspended 3–5 cm over the ground by a metal frame to allow free air circulation at the soil surface level to avoid an excessive heating. With these OTCs we aimed to apply a warming of 2–3°C, which is in line with several Atmosphere-Ocean General Circulation Models for the second half of this century in Central Spain (Giorgi and Lionello 2008; Stocker and others 2013).

Although precipitation predictions are subject to a higher level of uncertainty in the Mediterranean Basin, projected changes point to an intensification of water scarcity in this region (Giorgi and Lionello 2008; Stocker and others 2013). Thus, passive rainfall shelters based on the design of Yahdjian and Sala (2002) were used to achieve a reduction of rainfall amount of ~30% without changing the frequency of rainfall events. Each RS had an area of 1.68 m^2^ (1.4 × 1.2 m) and a roof composed by six gutters of methacrylate with an inclination of nearly 20° and a mean height of 1 m. The effectiveness of the passive rainfall shelters used have been previously tested (Escolar and others 2012; Maestre and others 2013; Ladrón de Guevara and others 2014).

### 2.5. Environmental monitoring

Rainfall and solar radiation were monitored by an on-site meteorological station (Onset Corp., Bourne, MA, USA). Air temperature and relative humidity in the different treatments were monitored with automated sensors (HOBO^®^ U23 Pro v2 Temp/RH Onset Corp., Pocasset, MA, USA). Soil surface temperature (first 2 cm) and soil water content (0-5 cm) were continually monitored with TMC20-HD and EC-5 sensors, respectively (Onset Corp. and Decagon, Inc., Pullman, WA, USA).

### 2.6. Litter decomposition

Six litterbags per subplot and species were placed in the field, accounting for a total of 480 litterbags (2 species x 6 timepoints x 2 light treatments x 4 climate treatments x 5 replicates). Five litterbags per combination of treatments and species were randomly removed for dry mass determinations approximately at 3, 5, 7, 11, 15 and 29 months after deployment in the field. Recovered litterbags were put in individual paper bags and were taken to the laboratory, where the lichen-litter was carefully cleared of soil, fauna and other lichens species with a soft brush. The litter samples were then dried in an oven at 60° C for 48 h before weighting.

Litter decomposition rate was estimated as the difference of dry mass between the initial and successive litterbag collection dates. The decomposition constant (*k*) of the litter of each lichen species was determined for climatic and light treatments using the single exponential decay model of (Olson 1963):

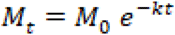

where *M*_t_ and *M*_0_ are the dry mass of the litter at time *t* and time 0. Thereby, *k* is the daily litter decay rate, which is commonly used in other experiments of litter decomposition (Lang and others 2009; Campbell and others 2010; King and others 2012; Almagro and others 2017).

### 2.7. Determinations of lichen mass chemistry

The contents of organic carbon (OC) and total nitrogen (TN) in the litter were determined at months 0, 11, and 29 after the setup of the experiment in five replicates per combination of treatments. Organic C was analysed with a colorimetric method after oxidation with a mixture of potassium dichromate and sulfuric acid (Anderson and Ingram 1994). Total N was determined with a Skalar Nutrient Autoanalyzer model SAN ++ (NUTILAB, URJC) after the sample digestion according to the Kjeldahl method, wherein a Cu + KSO_4_ catalyst is added to the sample and N-organic is oxidised to NH3-N by a digestion with H2SO_4_ (Radojevic and Bashkin 1999).

### 2.8. Statistical analyses

Differences in initial chemical composition (OC_0_, TN_0_ and C:N_0_) between species were analysed with Student’s t-tests. Also, we conducted statistical analyses on the remaining biomass of lichens, on the differences in their chemical composition 11 and 29 months after the beginning of the experiment and on the decomposition rates obtained. We analysed results at 11 and 29 months from experiment initialisation because mechanisms of litter degradation change throughout the decomposition process (García-Palacios and others 2016). We conducted three-way ANOVA’s to evaluate the effects of climate manipulations (C, RE, W and REW), light treatment (UV-block and UV-pass) and lichen species (*D. diacapsis* and *C. convoluta*) on remaining litter biomass (n = 5), decomposition rates (n = 5) and litter chemical composition (n=3). These factors were considered as fixed factors in these analyses. We added block as a random factor to control for microsite-specific differences among plots. To evaluate differences between levels within each fixed factor, Fisher’s least significant difference (LSD) mean comparison test was applied. Data were tested for ANOVA assumptions before analyses; *k* (day^−1^) values were log10-transformed to fulfil ANOVA assumptions. All statistical analyses were performed using SPSS 18.0 (SPSS Inc., Chicago, IL, USA).

## 3. Results

### 3.1. Effect of climate manipulation on environmental conditions

Accumulated rainfall during the experiment was 690 mm: 249, 386 and 55 mm for the first, second years and the last five months, respectively (Figure S4a). Mean monthly maximum air and soil temperatures (Figure S4b) were reached in July and August of both years (27°C and 32.5°C, respectively). On average, warming increased air temperature by 2.2°C with respect to the control, with maximum effect over the summer when differences reached up to 3.2°C. However, in the WRE treatment, the average increase with respect to the control was 1.4°C (Figure S5). Annual average relative humidity was 62% with autumns and summers having the highest (80%) and lowest (40%) values, respectively (Figure S5). On average, relative humidity was lower in the W and WRE treatments than in the control treatment by 4% and 3%, respectively. Variations in soil moisture closely followed rainfall events. Mean values during the experiment period were: 6.8% for C, 7.7% for RE, and 4% for W and WRE treatments.

### 3.2. Treatment effects on litter decomposition dynamics

After one year, the litter mass loss of both *D. diacapsis* and *C. Convoluta* was over 20% (Table S1, Figure 1). No significant effects of the treatments evaluated were observed on litter mass loss at this time (*P* > 0.05; Table S2). Nearly two and a half years after the experiment began, *D. diacapsis* lost almost 20% more mass than *C. convoluta* (*P* = 0.008). These effects were, however, modulated by the amount of UV radiation received, as indicated by a significant species × light interaction (*P* = 0.027; Figure 2). Thus, differences between species were only detected in the UV-pass treatment (*P* = 0.001; Table S3), and only *C. convoluta* had differences in remaining litter mass (*P* = 0.007) under the UV-pass treatment (Table S3; Figure 1). It must be noted, however, that the species × light × climate and species × climate interactions had P values of 0.082 (Table S2). Indeed, warming treatments (W and WRE) resulted in lower mass remaining values than no warming treatments (control and RE) but this effect was only significant for *D. diacapsis* in the UV-pass treatment (F = 3.2, *P* = 0.032, Figure 2).

**Figure 1.**
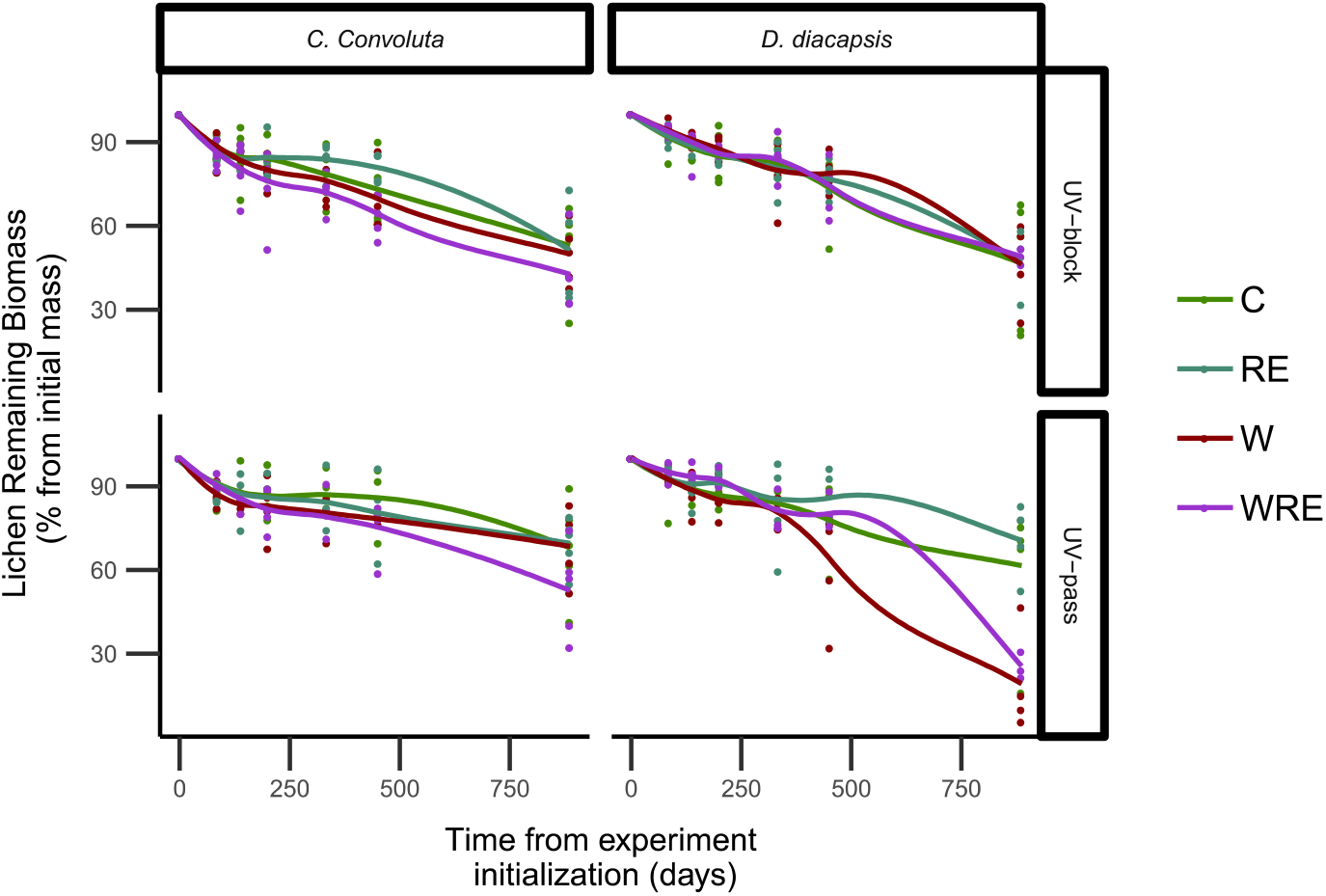
Lichen decomposition dynamics for *Diploschistes diacapsis* and *Cladonia convoluta* (columns) under two UV treatments (rows) and different climatic conditions. Lines are smoothed trends of the data. RE: rainfall exclusion, W: warming, WRE: warming and exclusion combination and C: control.

**Figure 2.**
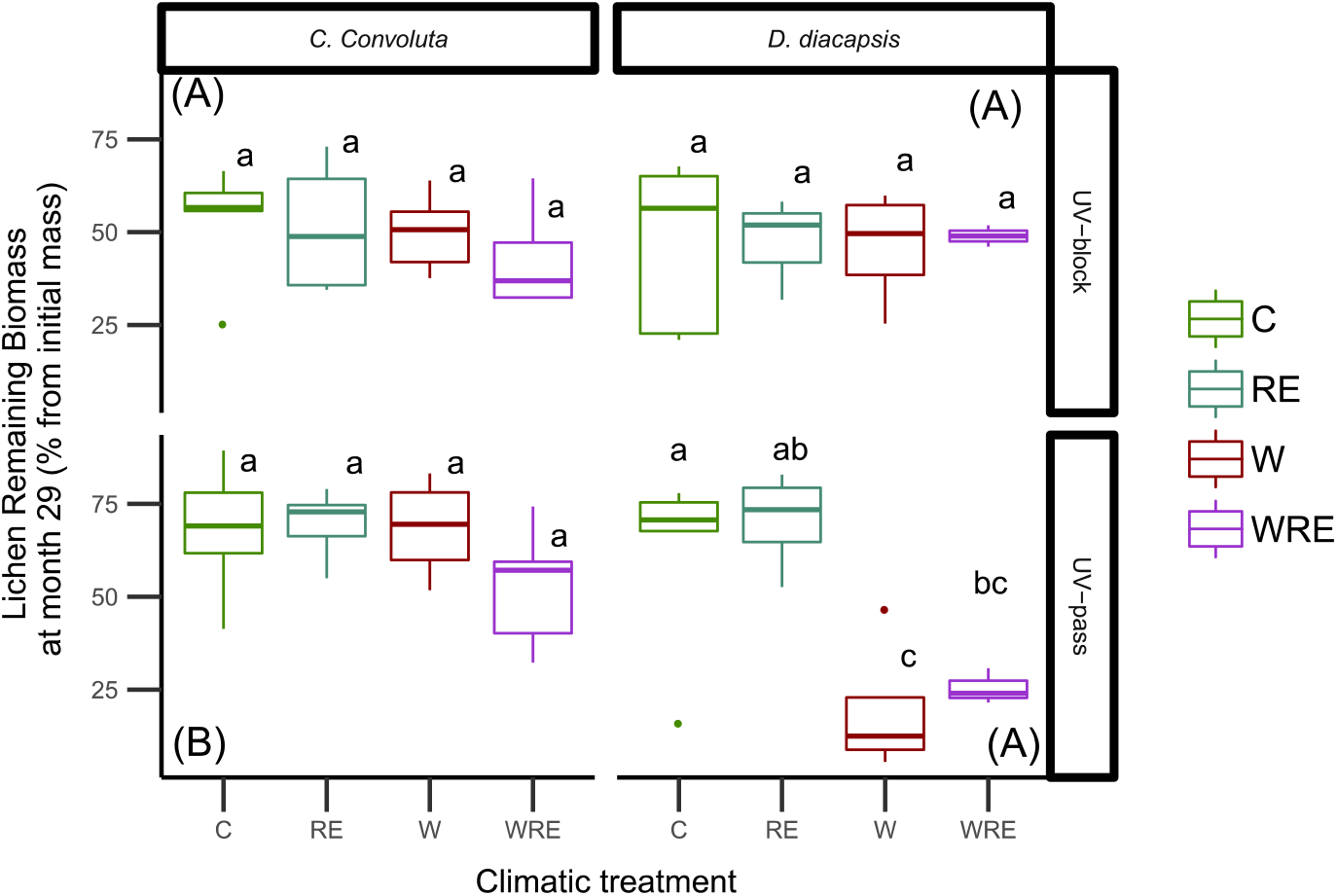
Mass remaining of *Diploschistes diacapsis* and *Cladonia convoluta* litter under different climate change and UV treatments 2.5 years after experimental setup. Different lowercase letters represent statistically significant differences (*P* < 0.05) when climatic treatments are compared within each panel. Uppercase letters in brackets represent homogeneous groups when comparing the mean of remaining biomass between each of the four panels. RE: rainfall exclusion, W: warming, WRE: warming and exclusion combination and C: control. Data are means ± SE (n = 5).

By the end of the experiment, the decomposition rate of *D. diacapsis* was higher (*k* = 0.0009 %/day) than that of *C. convoluta (k* = 0.0006 %/day). Significant three-way (species × light × climate) and two-way (species × light) interactions were observed when analysing litter decomposition rates (Table S2). Separate ANOVAS conducted for each species revealed that W and WRE accelerated litter decomposition rate in *D. diacapsis* and *C. convoluta*, respectively (Figure 3). Receiving more UV radiation favoured litter decay rate for *D. diacapsis* respect to *C. convoluta (F* = 12.24, *P* = 0.001). This effect was more important under warming treatments, where UV exposition increased the decomposition rates in *D. diacapsis* when compared to the UV-block treatment, while showing the opposite effect in *C. convoluta* (Figure 3). Also, *C. convoluta* had in general a higher *k* (day^−1^) in the UV-block treatment than in the UV-pass treatment (*F* = 7.379, *P* = 0.008). Contrasting effects where shown between light treatments for *D. diacapsis* depending on the climate treatment (Figure 3), finally rendering no significant differences overall (*F* = 0.719, *P* = 0.400). There were no differences between species in the absence of UV exposure (*F* = 0.004, *P* = 0.947).

**Figure 3.**
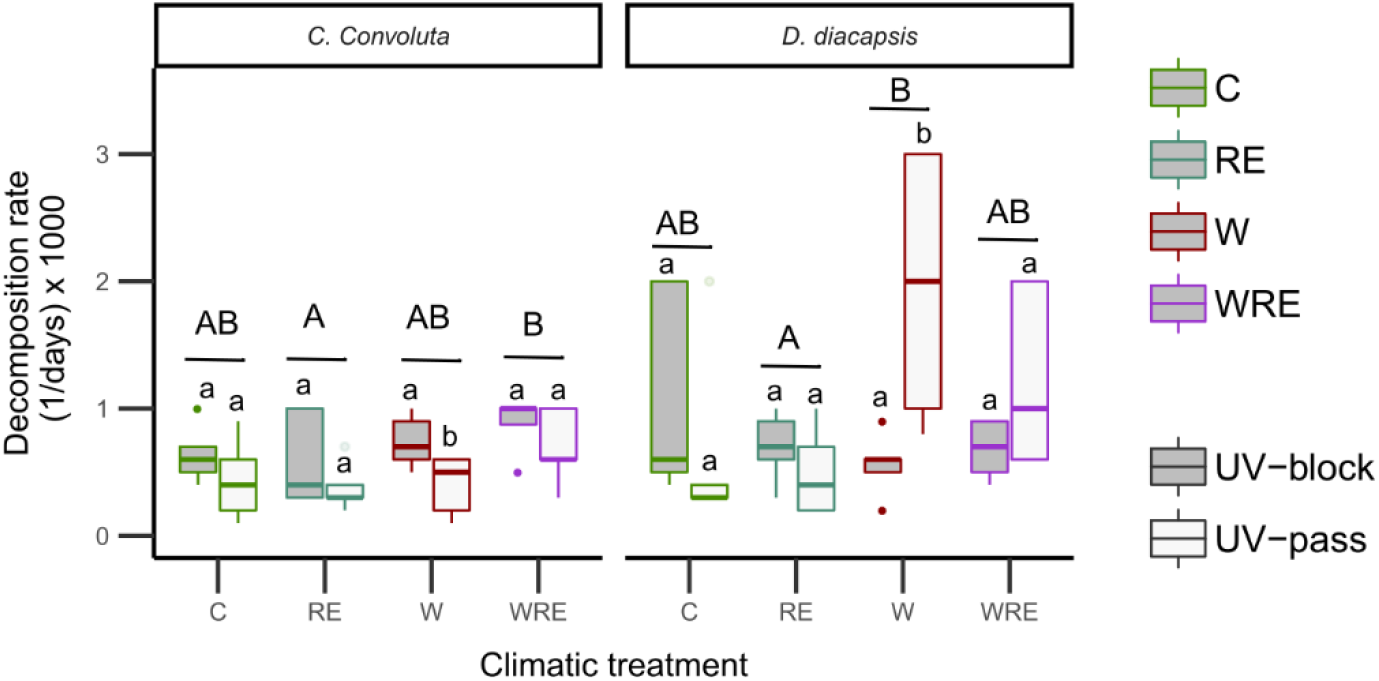
Decomposition rate of *Diploschistes diacapsis* and *Cladonia convoluta* litter under different climate change and UV treatments 2.5 years after experimental setup. Rest of legend as in Figure 2.

### 3.3. Treatment effects on carbon and nitrogen dynamics

The litter of *C. convoluta* had higher values of carbon and nitrogen content, and C:N ratio than that of *D. diacapsis* at the beginning of the experiment (Table S4). Differences in C and C:N ratio between species were maintained throughout the experiment regardless of the treatments, whereas N concentration values remained similar for both species throughout the experiment. In general, the light treatment was only significant after 29 months of experiment, promoting overall increases in the C:N ratio respect to UV-block treatments (Fig. 4). This effect was consistent among species (species × light interactions were not significant). However, depending on the species, different climatic treatments modulated C and N concentrations differently throughout the experiment (Table S5).

**Figure 4.**
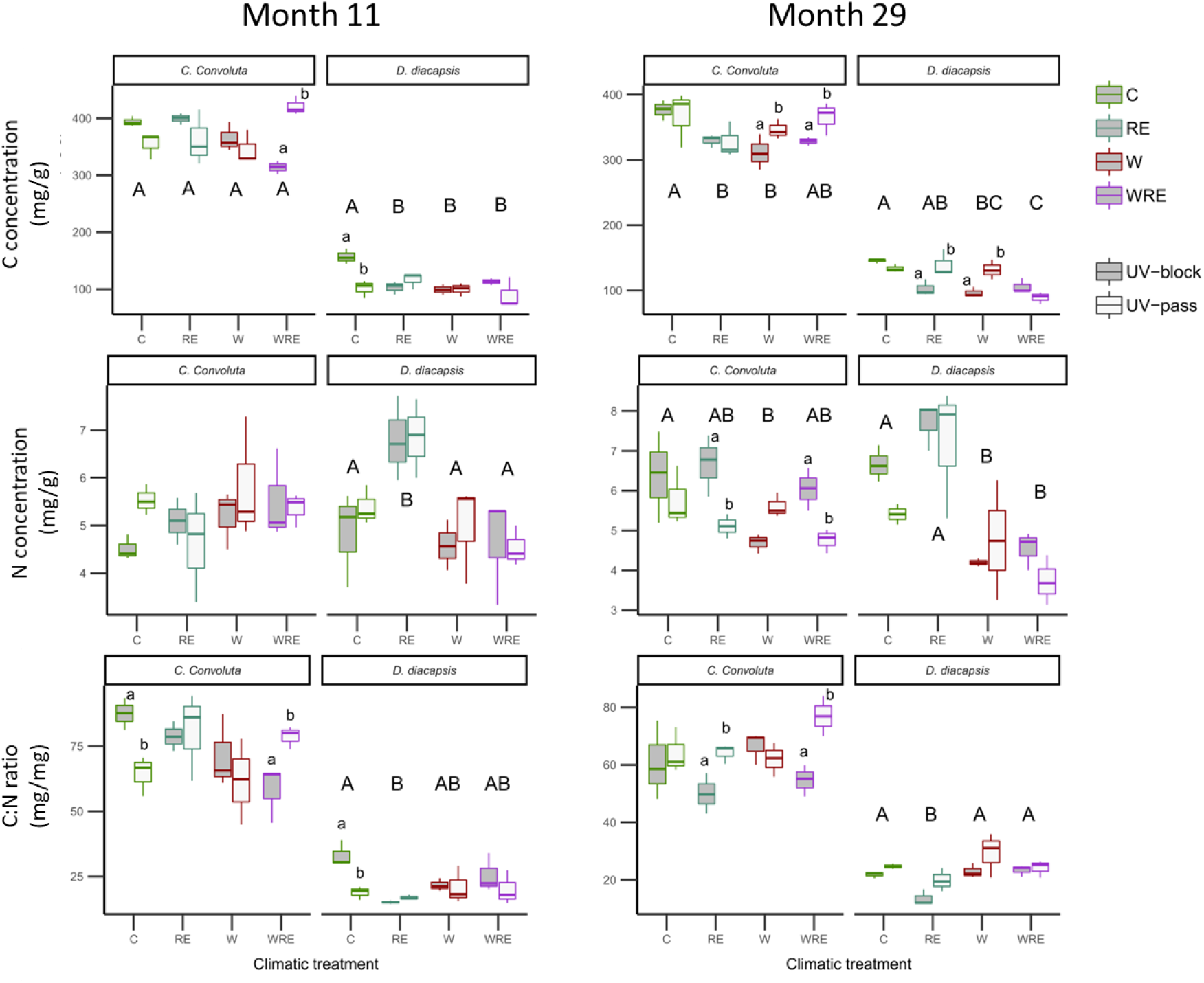
Carbon (top), Nitrogen (middle) and C:N ratio (bottom) concentrations at 11 (left) and 29 (right) months after the beginning of the experiment. Data are means ± SE (n = 3). Rest of legend as in Fig. 2.

*Diploschistes diacapsis* showed mainly effects of climatic treatments separately from light treatment effects. Some exception are the interactions found in C tissue concentration with the light treatment, which significantly increased this concentration in the RE and W treatments at month 29, and decreased that in the control treatment at month 11. In general, W and WRE treatments decreased both C and N concentration respect to control and RE treatments (Figure 4). This effect was clearer at month 29 than at month 11, where N concentration in the control treatment was similar to that in the warming treatments. Only the RE treatment exhibited lower C:N ratios than the rest of treatments; these were mainly driven by a higher N concentration respect to other treatments, which was also clear at month 11 (Figure 4).

In *C. convoluta*, the effects of climate treatments were only significant at month 29, with significant decreases of C and N concentrations under RE and W treatments respect to the control treatment (Figure 4). These differences, however were modulated by the light treatment. In rainfall treatments (RE and WRE), UV exposition significantly decreased N concentration while warming (W and WRE treatments) significantly increased C concentration respect to UV-block treatments. Most of these effects cancelled each other when evaluating the C:N ratio, which did not show clear effects of climate treatments. In the WRE and RE treatments, however, UV-pass treatments significantly increased C:N ratios respect to UV-block treatments (Figure 4).

## 4. Discussion

### 4.1. Litter decomposition rates of lichen species

Overall, litter decomposition rates of the lichens studied were low (*k* y^−1^ between 0.219 and 0.329). These values are comparable to the litter decomposition rates of some recalcitrant plant species in the Mediterranean region (e.g., *Bouteloua gracilis* and *Retama Sphaerocarpa, k~*_0_.36, see Brandt and others 2010; Almagro and others 2017; or *Q. coccifera* in southwest Spain, *k*~0.22, see Gallardo and Merino 1993) but still are faster than the most recalcitrant grass species present in the study area (*Stipa tenacissima, k* ~ 0.06, Almagro et al. 2015) or other species living in the same community (*H. Squamatum, k*~0.15, Prieto et al. 2019). Compared to other lichen species reported mostly in subarctic areas or coniferous forests, the decomposition rates shown here were lower than those of species of the same genus (*C. convoluta* vs. *C. stellaris k*~0.28, mass remaining ~ 40%) or morphology (*D. diacapsis* vs. *Umbilicaria hyperborea k* ~ 0.69, mass remaining ~ 20%, Lang and others 2009). This is probably related to site differences in soil and climate, which can limit litter decomposition in plants (Gallardo and Merino 1993). This is further supported by the fact that, although exhibiting lower C:N ratio compared to arctic communities (which indicates higher decomposability, Aber and others 1990), lichens in our experiment still decomposed slower than lichens of arctic communities (Lang and others 2009).

Litter decomposition is tightly linked to litter quality, being faster the lower the C:N ratio (Aber and others 1990). This premise explains why *D. diacapsis* (with C:N ratio ~ 30) decomposed faster than *C. convoluta* (C:N ratio ~ 56) in our experiment. The C:N ratio also controls the mobilization of N into the soil, as N usually is immobilised by microbial decomposers until C:N ratio ~ 20 (Manzoni and others 2010). Indeed, our results on the C:N ratios of *D.diacapsis* at the end of the experiment (~20) suggest that this species reached a stage in which N starts mineralisation while *C. convoluta* would still immobilise most of the nitrogen. This can also explain differences in N content of both lichens at the end of the experiment and a faster litter decomposition rate of *D. dicapsis*, especially during the second year of the experiment, but specific measurements to assess the mineralised nitrogen should be taken to further corroborate this hypothesis.

Decomposition of lichen tissues was accelerated mainly in the second year of our experiment. This concurs with temporal patterns of decomposition evidenced for other lichen species (Hagemann and Moroni 2015) and grasses (Wang and others 2017) in areas with periods of unfavourable conditions such as freezing. However, it contrasts with rapid losses of biomass at early stages that are more common in plant decomposition experiments in Mediterranean ecossytems (Almagro and others 2015). It is interesting to note that differences between lichen species were only apparent during this second year. According to other studies, legacy effects related to litter quality are more apparent on late stages of decomposition, when the relative importance of biotic processes linked to microarthropods and nematodes is reduced (García-Palacios and others 2016). Thus, our results also support the hypothesis that differences in litter decomposition rates between species is mainly controlled by litter quality.

Apart from the C:N ratio, other chemical components of litter may have a fundamental role in litter decomposition, particularly in lichens (Lang and others 2009). In *D. diacapsis*, almost 40% of lichenic organic carbon is labile (Miralles and others 2013). This labile carbon can be lost immediately by active microbial synthesis (Chantigny and others 2006) or by microbial respiration of CO_2_ via litter decomposition (Delgado-Baquerizo and others 2015b). Both studied species also differ in the content of lichenic substances; *C. convoluta* presents phenolic compounds like, fumarprotocetraric acid (aromatic aldehyde), usnic acid, zeorin (Triterpenoides) and psoromic acid (Pino-Bodas and others 2010), while *D. diacapsis* presents diploschistesic, lecanoric and or sellinic acid (Nash and others 2002). Usnic acid is associated with lichen thallus protection to UV radiation, while fumarprotocetraric acid had hydrophobic properties (Barreno and Párez-Ortega 2003). Both compounds have also high antimicrobial activity that inhibit bacteria and fungi. The protection provided by lichenic substances to alive lichens could affect the decomposition process of their tissues (Kosanić and others 2014). For instance, phenolic compounds have been shown to reduce the positive effects of microarthropods on the decomposition of lichen litter, although these substances reduce rapidly through decomposition after one year (Asplund and Wardle 2013).

### 4.2. Effect of climate manipulation on litter decomposition

According to previous results in other climate manipulation experiments carried out with plants (Almagro and others 2015), we expected warming and rainfall exclusion to decrease overall decomposition rates due to lower water availability under these treatments. However, warming increased litter decomposition rates of *D. diacapsis*, a response accompanied by higher N mobilization (as indicated by N concentration rates, Fig. 3), and of *C. Convoluta*, albeit in this species this response was only observed when warming was combined with reduced precipitation. Albeit initially unexpected, our results could explain the higher respiration rates observed under warming at our study site during the first years after the setup of the experiment (Maestre and others 2013). Furthermore, our results indicate that these increases in soil respiration with warming are not only due to a higher lichen mortality, but also to a higher decomposition of their tissues. Interestingly, this positive effect of warming on soil respiration disappears 8-10 years after the setup of the experiment (Dacal and others 2020), when the mortality of biocrust-forming lichens was mostly halted compared to the initial years of the experiment (Ladrón de Guevara and others 2018).

While we can only hypothesise the reasons for a higher decomposition under warming treatments, we speculate it could be driven by two alternative hypotheses. First, summer is an unfavorable period for biotic activity in the study area due to the acute water scarcity. In contrast, during winter water is no longer a limiting factor although low temperatures may reduce biotic activity significantly (despite the still warm temperature compared to other areas). Thus, the warming treatment could have accelerated decomposition processes during winter and early spring by creating a warmer micro environment prone to decomposition in the season when it rains more. Another explanation may be the existence of some kind of antimicrobial effect exerted by lichens (Kosanić and others 2014). The effect of soil humidity on litter decomposition rates is partly mediated by microbial activity, which degrades litter (Manzoni and others 2010; García-Palacios and others 2016). Thus, the antimicrobial compounds present in lichen tissues may have obscured the effect of soil humidity on their degradation by impeding efficient microbial activity. This is especially relevant for biocrust-forming lichens, which live in tight interaction with soil microorganisms and, unlike plants, developed a series of chemical compounds to compete with them (Kosanic and Ranković 2015).

Regardless of the mechanisms underlying them, our results indicate that abiotic drivers of decomposition would take a predominant role in lichen degradation. Among these drivers thermal degradation has been described to play a role in litter decomposition, especially in drylands (Brandt and others 2007; Gallo and others 2009; Almagro and others 2017). Our results support this idea, as evidenced from the effects of the light treatment observed (Figures 1 and 2).

### 4.3. Effects of UV radiation on litter decomposition

Observed increases in decomposition rates in the warming treatment were partly mediated by the effects of light treatment. Unlike with previous observations carried out with plant litter (Brandt and others 2010; King and others 2012; Almagro and others 2017; Wang and others 2017), UV exposure only accelerated lichen litter decomposition in conjunction with warming in *D. diacapsis*. However, *C. convoluta* litter showed a contrasting behavior and tended to slow down the decomposition process with the warming when exposed to UV. Photodegradation is an increasingly recognized driver of litter decomposition in arid lands (Austin and Vivanco 2006; King and others 2012), and may increase degradation by mechanic fragmentation of lichens and thermal degradation (Gallo and others 2006; Day and others 2007). On the other hand, increased UV radiation can slow down the decomposition process due to the negative effect on microbial and arthropod life forms (Brandt and others 2007, 2010). Thus, the effect of photodegradation on litter decomposition is the result of a balance between abiotic and biotic processes.

The chemical composition and morphology of the lichen thalli may be an important driver explaining the contrasting decomposition behavior of the two lichen species studied. For instance, differences in content, number and complexity of carotenoids between some species of *D. diacapsis* and *C. convoluta* have been reported (e.g. *D. muscorum* and *C. furcaiu*, Czeczuga and others 1995), which is also apparent by their different color. A higher content of pigments in *C. Convoluta* or other compounds that absorb UV radiation (e.g., usnic acid) may decrease the efficiency of direct photodegradation when compared with *D. diacapsis*, which is white and show less usnic acid and different composition of carotenoids than *C. convoluta* (Nash and others 2002). Similarly, *D. diacapsis* is more exposed to radiation due to its plain morphology, whereas *C. convoluta* is a foliose lichen and its higher structural complexity and volume may produce more diffuse impact of radiation in the thallus. These drivers have been previously overlooked in plants, which exhibit less variation in their morphology and whose composition in UV absorbing pigments is more homogeneous than in lichens. Future studies need to test these hypotheses by examining the importance of photodegradation in a wider variety of lichens respect to its pigment content.

Apart of the overall decomposition rates, photodegradation played a predominant role on the modulation C and N concentration of lichen species studied. Our results suggest that these effects are more apparent in *C. convoluta*, the species with the lowest decomposition rates. It is interesting to note that UV exposure effects were more apparent in the climate change treatments. In particular, the lower N concentration rates found in RE and WRE treatments suggests that UV radiation may even mobilize N regardless of the C:N ratio of the thallus, thus increasing abiotic decomposition (Austin and Vivanco 2006). This effect is probably more important in dry lichens, which would explain why this result was significant only in the RE and WRE treatments. An important point is that this N mobilization by UV radiation prevents the C:N ratios to decrease, and thus may suppose an impediment to microbes for mobilizing N compounds through the decomposition process. Our results also suggest that the role of photodegradation will be more important with ongoing climate change.

## 5. Concluding remarks

To the best of our knowledge, this is the first study assessing the decomposition of soil lichens under simulated climate change in drylands. Our results showed that dryland lichens may decompose faster than plants growing in the same area and under the same climatic conditions, but slower than lichens from temperate and arctic ecosystems. Altogether, our results show that the degradation of lichen litter is largely controlled by abiotic factors (UV radiation) and the chemical composition of lichen thalli. An interesting contribution of our study is that the effects of climate change on lichen decomposition may interact importantly with the abiotic and biotic drivers of the decomposition process, highlighting the importance of further investigating the dynamics and drivers of lichen decomposition to fully understand the impacts of climate change on soil carbon and nutrient dynamics in drylands. Future studies should focus on biotic drivers of lichenic decomposition to fully understand the contrasting results obtained here and nuance the particularities of lichen decomposition in these environments. They should also take into account the composition of secondary components of lichens (such as antimicrobial peptides and phenolic compounds) as a relevant driver of lichen decomposition. Our findings also stress the importance of considering biocrusts and the decomposition of their tissues when modelling carbon cycling responses to climate change in drylands.

## Acknowledgements

This research was supported by the European Research Council [ERC grant nos. 242658 (BIOCOM) and 647038 (BIODESERT) awarded to F.T.M.]. M.B. acknowledges support from a Juan de la Cierva Formación grant from the Spanish Ministry of Economy and Competitiveness (FJCI-2018-036520-I). F.T.M. acknowledges support from Generalitat Valenciana (CIDEGENT/2018/041), the Alexander von Humboldt Foundation, and the Synthesis Centre for Biodiversity Sciences (sDiv) of the German Centre for Integrative Biodiversity Research (iDiv). Author contributions: M.B., A.R. and F.T.M designed the experiment. V.O., B.G., L.G. and D.M. monitored the experiment and took laboratory and weight measurements. A.R., M.B. and D.O.M. performed statistical analyses. M.B. and D.O.M. wrote the paper and all authors contributed to further editions and corrections. Data and materials are available from figshare (https://figshare.com/s/d5fa2310717821c6b779; see (Berdugo and others 2020)).

## References

Aber JD, Melillo JM, McClaugherty CA. 1990. Predicting long-term patterns of mass loss, nitrogen dynamics, and soil organic matter formation from initial fine litter chemistry in temperate forest ecosystems. Can J Bot 68:2201–8. https://doi.org/10.1139/b90-287

Almagro M, Maestre FT, Martínez-López J, Valencia E, Rey A. 2015. Climate change may reduce litter decomposition while enhancing the contribution of photodegradation in dry perennial Mediterranean grasslands. Soil Biol Biochem 90:214–23. http://www.sciencedirect.com/science/article/pii/S0038071715002758

Almagro M, Martínez-López J, Maestre FT, Rey A. 2017. The Contribution of Photodegradation to Litter Decomposition in Semiarid Mediterranean Grasslands Depends on its Interaction with Local Humidity Conditions, Litter Quality and Position. Ecosystems 20:527–42. https://doi.org/10.1007/s10021-016-0036-5

Anderson JM, Ingram JSI. 1994. Tropical soil biology and fertility: A handbook of methods. Soil Sci 157:265.

Asplund J, Wardle DA. 2013. The impact of secondary compounds and functional characteristics on lichen palatability and decomposition. J Ecol 101:689–700. https://doi.org/10.1111/1365-2745.12075

Austin AT, Vivanco L. 2006. Plant litter decomposition in a semi-arid ecosystem controlled by photodegradation. Nature 442:555–8. https://doi.org/10.1038/nature05038

Barreno RE, Párez-Ortega S. 2003. Biología de los líquenes. In: Barreno RE, Pérez-Ortega S, editors. Líquenes de la Reserva Natural Integral de Muniellos, Asturias. Oviedo, Spain: Conserjería de Medio Ambiente, Ordenación del Territorio e Infraestructuras del Principado de Asturias. pp 65–82.

Belnap J, Lange OL. 2013. Biological Soil Crusts: Structure, Function and Management. 2nd ed. (Belnap J, Lange OL, editors.). Berlin, Heidelberg: Springer, Berlin, Heidelberg

Berdugo M, Mendoza-Aguilar DO, Rey A, Ochoa V, Gozalo B, García-Huss L, Maestre FT. 2020. Dataset form the study: Litter decomposition rates of biocrust-forming lichens are similar to that of vascular plants and are affected by warming in semiarid grasslands. figshare. doi: 10.6084/m9.figshare.11822538.

Bowker MA, Mau RL, Maestre FT, Escolar C, Castillo-Monroy AP. 2011. Functional profiles reveal unique ecological roles of various biological soil crust organisms. Funct Ecol 25:787–95. https://doi.org/10.1111/j.1365-2435.2011.01835.x

Brandt LA, King JY, Hobbie SE, Milchunas DG, Sinsabaugh RL. 2010. The Role of Photodegradation in Surface Litter Decomposition Across a Grassland Ecosystem Precipitation Gradient. Ecosystems 13:765–81. https://doi.org/10.1007/s10021-010-9353-2

Brandt LA, King JY, Milchunas DG. 2007. Effects of ultraviolet radiation on litter decomposition depend on precipitation and litter chemistry in a shortgrass steppe ecosystem. Glob Chang Biol 13:2193–205. https://doi.org/10.1111/j.1365-2486.2007.01428.x

Caldiz MS, Brunet J, Nihlgård B. 2007. Lichen litter decomposition in *Nothofagus* forest of northern Patagonia: biomass and chemical changes over time. Bryologist 110:266–73. https://doi.org/10.1639/0007-2745(2007)110[266:LLDINF]2.0.CO

Campbell J, Fredeen AL, Prescott CE. 2010. Decomposition and nutrient release from four epiphytic lichen litters in sub-boreal spruce forests. Can J For Res 40:1473–84. https://doi.org/10.1139/X10-071

Cano-Díaz C, Mateo P, Muñoz-Martín MÁ, Maestre FT. 2018. Diversity of biocrust-forming cyanobacteria in a semiarid gypsiferous site from Central Spain. J Arid Environ 151:83–9. http://www.sciencedirect.com/science/article/pii/S0140196317302148

Castillo-Monroy AP, Maestre FT, Rey A, Soliveres S, García-Palacios P. 2011. Biological Soil Crust Microsites Are the Main Contributor to Soil Respiration in a Semiarid Ecosystem. Ecosystems 14:835–47. http://www.springerlink.com/index/10.1007/s10021-011-9449-3. Last accessed 14/03/2013

Chantigny MH, Angers DA, Kaiser K, Kalbitz K. 2006. Extraction and Characterization of Disolved Organic Matter. In: Carter MR, Gregorich EG, editors. Soil Sampling and Methods of Analysis. Second. Canadian Society of Soil Science. pp 617–35.

Cherlet M, Hutchinson C, Reynolds J, Hill J, Sommer S, von Maltitz G. 2018. World Atlas of Desertification. 3rd ed. (Cherlet M, Hutchinson C, Reynolds J, Hill J, Sommer S, von Maltitz G, editors.). Luxembourg: Publication Offise of the European Union

Ciais P, Bombelli A, Williams M, Piao S L, Chave J, Ryan C M, Henry M, Brender P, Valentini R. 2011. The carbon balance of Africa: synthesis of recent research studies. Philos Trans R Soc A Math Phys Eng Sci 369:2038–57. https://doi.org/10.1098/rsta.2010.0328

Czeczuga B, Von Arb C, Lumbsch HT. 1995. Carotenoids in lichens from the Swiss Alps. FEDDES Repert 106:173–8.

Dacal M, García-Palacios P, Asensio S, Gozalo B, Ochoa V, Maestre FT. 2020. Abiotic and biotic drivers underly short- and long-term soil respiration responses to experimental warming in a dryland ecosystem. bioRxiv:2020.01.13.903880. http://biorxiv.org/content/early/2020/01/15/2020.01.13.903880.abstract

Darrouzet-Nardi A, Reed SC, Grote EE, Belnap J. 2015. Observations of net soil exchange of CO2 in a dryland show experimental warming increases carbon losses in biocrust soils. Biogeochemistry 126:363–78.

Day TA, Zhang ET, Ruhland CT. 2007. Exposure to solar UV-B radiation accelerates mass and lignin loss of Larrea tridentata litter in the Sonoran Desert. Plant Ecol 193:185–94. https://doi.org/10.1007/s11258-006-9257-6

Delgado-Baquerizo M, Gallardo A, Covelo F, Prado-Comesaña A, Ochoa V, Maestre FT. 2015a. Differences in thallus chemistry are related to species-specific effects of biocrust-forming lichens on soil nutrients and microbial communities. Funct Ecol 29:1087–98. https://doi.org/10.1111/1365-2435.12403

Delgado-Baquerizo M, García-Palacios P, Milla R, Gallardo A, Maestre FT. 2015b. Soil characteristics determine soil carbon and nitrogen availability during leaf litter decomposition regardless of litter quality. Soil Biol Biochem 81:134–42. http://www.sciencedirect.com/science/article/pii/S0038071714003873

Dirks I, Navon Y, Kanas D, Dumbur R, Grünzweig JM. 2010. Atmospheric water vapor as driver of litter decomposition in Mediterranean shrubland and grassland during rainless seasons. Glob Chang Biol 16:2799–812. https://doi.org/10.1111/j.1365-2486.2010.02172.x

Escolar C, Maestre FT, Rey A. 2015. Biocrusts modulate warming and rainfall exclusion effects on soil respiration in a semi-arid grassland. Soil Biol Biochem 80:9–17.

Escolar C, Martínez I, Bowker MA, Maestre FT. 2012. Warming reduces the growth and diversity of biological soil crusts in a semi-arid environment: implications for ecosystem structure and functioning. Philos Trans R Soc London B Biol Sci 367:3087–99. http://www.ncbi.nlm.nih.gov/pubmed/23045707. Last accessed 06/03/2013

Ferrenberg S, Reed SC, Belnap J. 2015. Climate change and physical disturbance cause similar community shifts in biological soil crusts. Proc Natl Acad Sci 112:12116–21.

Gallardo A, Merino J. 1993. Leaf Decomposition in Two Mediterranean Ecosystems of Southwest Spain: Influence of Substrate Quality. Ecology 74:152–61. https://doi.org/10.2307/1939510

Gallo ME, Porras-Alfaro A, Odenbach KJ, Sinsabaugh RL. 2009. Photoacceleration of plant litter decomposition in an arid environment. Soil Biol Biochem 41:1433–41. http://www.sciencedirect.com/science/article/pii/S0038071709001308

Gallo ME, Sinsabaugh RL, Cabaniss SE. 2006. The role of ultraviolet radiation in litter decomposition in arid ecosystems. Appl Soil Ecol 34:82–91. http://www.sciencedirect.com/science/article/pii/S0929139306000187

García-Palacios P, Escolar C, Dacal M, Delgado-Baquerizo M, Gozalo B, Ochoa V, Maestre FT. 2018. Pathways regulating decreased soil respiration with warming in a biocrust-dominated dryland. Glob Chang Biol 24:4645–56. https://doi.org/10.1111/gcb.14399

García-Palacios P, Maestre FT, Kattge J, Wall DH. 2013. Climate and litter quality differently modulate the effects of soil fauna on litter decomposition across biomes. Ecol Lett 16:1045–53. https://doi.org/10.1111/ele.12137

García-Palacios P, Shaw EA, Wall DH, Hättenschwiler S. 2016. Temporal dynamics of biotic and abiotic drivers of litter decomposition. Ecol Lett 19:554–63. https://doi.org/10.1111/ele.12590

Giorgi F, Lionello P. 2008. Climate change projections for the Mediterranean region. Glob Planet Change 63:90–104. http://www.sciencedirect.com/science/article/pii/S0921818107001750

Hagemann U, Moroni MT. 2015. Moss and lichen decomposition in old-growth and harvested high-boreal forests estimated using the litterbag and minicontainer methods. Soil Biol Biochem 87:10–24. http://www.sciencedirect.com/science/article/pii/S0038071715001443

Huang J, Yu H, Guan X, Wang G, Guo R. 2015. Accelerated dryland expansion under climate change. Nat Clim Chang 6:166–71.

IUSS Working Group WRB. 2015. World Reference Base for Soil Resources 2014, update 2015. Rome

King JY, Brandt LA, Adair EC. 2012. Shedding light on plant litter decomposition: advances, implications and new directions in understanding the role of photodegradation. Biogeochemistry 111:57–81. https://doi.org/10.1007/s10533-012-9737-9

Kosanic M, Ranković B. 2015. Chapter 3: Lichen Secondary Metabolites as Potential Antibiotic Agents. In: Rankovic B, editor. Lichen secondary metabolites. Bioactive properties and pharmaceutical potential. Springer International Publishing Switzerland. pp 91–104.

Kosanić M, Ranković B, Stanojković T, Rančić A, Manojlović N. 2014. Cladonia lichens and their major metabolites as possible natural antioxidant, antimicrobial and anticancer agents. LWT - Food Sci Technol 59:518–25. http://www.sciencedirect.com/science/article/pii/S0023643814002527

Ladrón de Guevara M, Gozalo B, Raggio J, Lafuente A, Prieto M, Maestre FT. 2018. Warming reduces the cover, richness and evenness of lichen-dominated biocrusts but promotes moss growth: insights from an 8 yr experiment. New Phytol 220:811–23. https://doi.org/10.1111/nph.15000

Ladrón de Guevara M, Lázaro R, Quero JLJL, Ochoa V, Gozalo B, Berdugo M, Uclés O, Escolar C, Maestre FTFT. 2014. Simulated climate change reduced the capacity of lichen-dominated biocrusts to act as carbon sinks in two semi-arid Mediterranean ecosystems. Biodivers Conserv 23:1787–807.

Lang SI, Cornelissen JHC, Klahn T, Van Logtestijn RSP, Broekman R, Schweikert W, Aerts R. 2009. An experimental comparison of chemical traits and litter decomposition rates in a diverse range of subarctic bryophyte, lichen and vascular plant species. J Ecol 97:886–900. https://doi.org/10.1111/j.1365-2745.2009.01538.x

Li S, Liu W-Y, Li D-W, Li Z-X, Song L, Chen K, Fu Y. 2014. Slower rates of litter decomposition of dominant epiphytes in the canopy than on the forest floor in a subtropical montane forest, southwest China. Soil Biol Biochem 70:211–20. http://www.sciencedirect.com/science/article/pii/S0038071713004756

Maestre FT, Delgado-Baquerizo M, Jeffries TC, Eldridge DJ, Ochoa V, Gozalo B, Quero JL, García-Gómez M, Gallardo A, Ulrich W, Bowker MA, Arredondo T, Barraza C, Bran D, Florentino A, Gaitán J, Gutiérrez JR, Huber-Sannwald E, Jankju M, Mau RL, Miriti M, Naseri K, Ospina A, Stavi I, Wang D, Woods NN, Yuan X, Zaady E, Singh BK. 2015. Increasing aridity reduces soil microbial diversity and abundance in global drylands. Proc Natl Acad Sci 112:15684–9.

Maestre FTFT, Escolar C, de Guevara ML, Quero JLJL, Lázaro R, Delgado-Baquerizo M, Ochoa V, Berdugo M, Gozalo B, Gallardo A, Guevara ML, Quero JLJL, Lázaro R, Delgado-Baquerizo M, Ochoa V, Berdugo M, Gozalo B, Gallardo A, de Guevara ML, Quero JLJL, Lázaro R, Delgado-Baquerizo M, Ochoa V, Berdugo M, Gozalo B, Gallardo A. 2013. Changes in biocrust cover drive carbon cycle responses to climate change in drylands. Glob Chang Biol 19:3835–47.

Manzoni S, Trofymow JA, Jackson RB, Porporato A. 2010. Stoichiometric controls on carbon, nitrogen, and phosphorus dynamics in decomposing litter. Ecol Monogr 80:89–106. https://doi.org/10.1890/09-0179.1

Maphangwa KW, Musil CF, Raitt L, Zedda L. 2012. Experimental climate warming decreases photosynthetic efficiency of lichens in an arid South African ecosystem. Oecologia 169:257–68.

McCulley RL, Burke IC, Nelson JA, Lauenroth WK, Knapp AK, Kelly EF. 2005. Regional Patterns in Carbon Cycling Across the Great Plains of North America. Ecosystems 8:106–21. https://doi.org/10.1007/s10021-004-0117-8

McCune B, Daly WJ. 1994. Consumption and Decomposition of Lichen Litter in a Temperate Coniferous Rainforest. Lichenol 26:67–71. https://www.cambridge.org/core/article/consumption-and-decomposition-of-lichen-litter-in-a-temperate-coniferous-rainforest/EA2E5647D592D106FDEC8C78320F57B5

Miralles I, Trasar-Cepeda C, Leirós MC, Gil-Sotres F. 2013. Labile carbon in biological soil crusts in the Tabernas desert, SE Spain. Soil Biol Biochem 58:1–8. http://www.sciencedirect.com/science/article/pii/S0038071712004488

Nash T., Ryan BD, Gries C, Bungatz F. 2002. Lichen flora of the Greater Sonoran Desert Region. Vol.1. (Nash T., Ryan BD, Gries C, Bungatz F, editors.). Tempe, Arizona: Arizona State University https://lichenportal.org/cnalh/taxa/index.php?taxon=52882&clid=1130

Olson JS. 1963. Energy Storage and the Balance of Producers and Decomposers in Ecological Systems. Ecology 44:322–31. https://doi.org/10.2307/1932179

Pino-Bodas R, Martín MP, Burgaz ANAR. 2010. Insight into the Cladonia convoluta-C. foliacea (Cladoniaceae, Ascomycota) complex and related species, revealed through morphological, biochemical and phylogenetic analyses. Syst Biodivers 8:575–86. https://doi.org/10.1080/14772000.2010.532834

Poulter B, Frank D, Ciais P, Myneni RB, Andela N, Bi J, Broquet G, Canadell JG, Chevallier F, Liu YY, Running SW, Sitch S, van der Werf GR. 2014. Contribution of semi-arid ecosystems to interannual variability of the global carbon cycle. Nature 509:600. https://doi.org/10.1038/nature13376

Prieto I, Almagro M, Bastida F, Querejeta JI. 2019. Altered leaf litter quality exacerbates the negative impact of climate change on decomposition. J Ecol 0. https://doi.org/10.1111/1365-2745.13168

Radojevic M, Bashkin VN. 1999. Practical Environmental Analysis. 2nd ed. (Radojevic M, Bashkin VN, editors.). Cambridge: Royal Society of Chemistry, Cambridge, UK

Reed SC, Coe KK, Sparks JP, Housman DC, Zelikova TJ, Belnap J. 2012. Changes to dryland rainfall result in rapid moss mortality and altered soil fertility. Nat Clim Chang 2:752. https://doi.org/10.1038/nclimate1596

Resplandy L, Keeling RF, Rödenbeck C, Stephens BB, Khatiwala S, Rodgers KB, Long MC, Bopp L, Tans PP. 2018. Revision of global carbon fluxes based on a reassessment of oceanic and riverine carbon transport. Nat Geosci 11:504–9. https://doi.org/10.1038/s41561-018-0151-3

Rodriguez-Caballero E, Belnap J, Büdel B, Crutzen PJ, Andreae MO, Pöschl U, Weber B. 2018. Dryland photoautotrophic soil surface communities endangered by global change. Nat Geosci 11:185–9. https://doi.org/10.1038/s41561-018-0072-1

Rutledge S, Campbel DI, Baldocchi D, Schipper LA. 2010. Photodegradation leads to increased carbon dioxide losses from terrestrial organic matter. Glob Chang Biol 16:3065–74. https://doi.org/10.1111/j.1365-2486.2009.02149.x

Safriel UN, Adeel Z. 2005. Dryland Systems. In: Hassan R, Scholes R, Ash N, editors. Ecosystems and Human Well-being: Current State and Trends. Vol. 1. Washington D.C., USA: Island Press. pp 632–62.

Stocker TF, Qin D, Plattner GK, Tignor M, Allen SK, Boschung J, Nauels A, Xia Y, Bex B, Midgley BM. 2013. Climate change 2013: the physical science basis. Contribution of working group I to the fifth assessment report of the intergovernmental panel on climate change. Geneva: Cambridge University Press

Thomas AD. 2012. Impact of grazing intensity on seasonal variations in soil organic carbon and soil CO2 efflux in two semiarid grasslands in southern Botswana. Philos Trans R Soc B Biol Sci 367:3076–86. https://doi.org/10.1098/rstb.2012.0102

Throop HL, Archer SR. 2009. Resolving the Dryland Decomposition Conundrum: Some New Perspectives on Potential Drivers BT - Progress in Botany. In: Lüttge U, Beyschlag W, Büdel B, Francis D, editors. Berlin, Heidelberg: Springer Berlin Heidelberg. pp 171–94. https://doi.org/10.1007/978-3-540-68421-3_8

Wang J, Yang S, Zhang B, Liu W, Deng M, Chen S, Liu L. 2017. Temporal dynamics of ultraviolet radiation impacts on litter decomposition in a semi-arid ecosystem. Plant Soil 419:71–81. https://doi.org/10.1007/s11104-017-3290-1

Wardle DA, Bardgett RD, Klironomos JN, Setälä H, van der Putten WH, Wall DH. 2004. Ecological Linkages Between Aboveground and Belowground Biota. Science (80-) 304:1629 LP – 1633. http://science.sciencemag.org/content/304/5677/1629.abstract

Wetmore CM. 1982. Lichen Decomposition in a Black Spruce Bog. Lichenol 14:267–71. https://www.cambridge.org/core/article/lichen-decomposition-in-a-black-spruce-bog/EE6840AA1B5DE290E41B0B957F095E45

Yahdjian L, Sala OE. 2002. A rainout shelter design for intercepting different amounts of rainfall. Oecologia 133:95–101. https://doi.org/10.1007/s00442-002-1024-3

