## Supplementary Figures and Tables for "Litter decomposition rates of biocrust-forming lichens are similar to that of vascular plants and are affected by warming in a semiarid grassland"

**Supplementary material for the article: "Litter decomposition rates of biocrust-forming lichens are similar to that of vascular plants and are affected by warming in semiarid grasslands"**

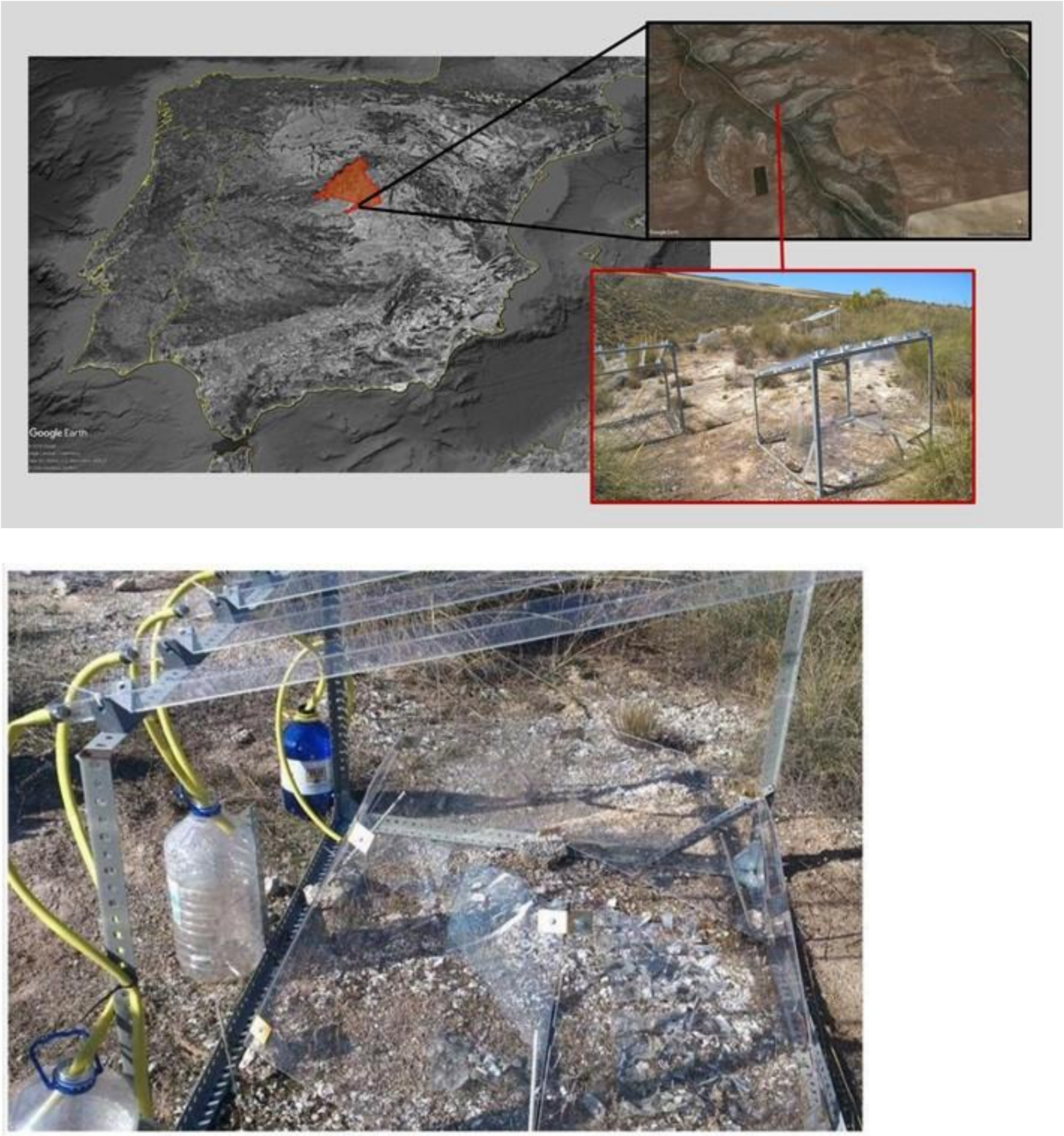


**Figure S1.** Location of the Aranjuez Experimental Station and view of the passive rainfall shelters and open top chamber for climate change experiment (upper panels), and detail of a warming and rainfall exclusion experimental plot (low picture; photo by Laura García).

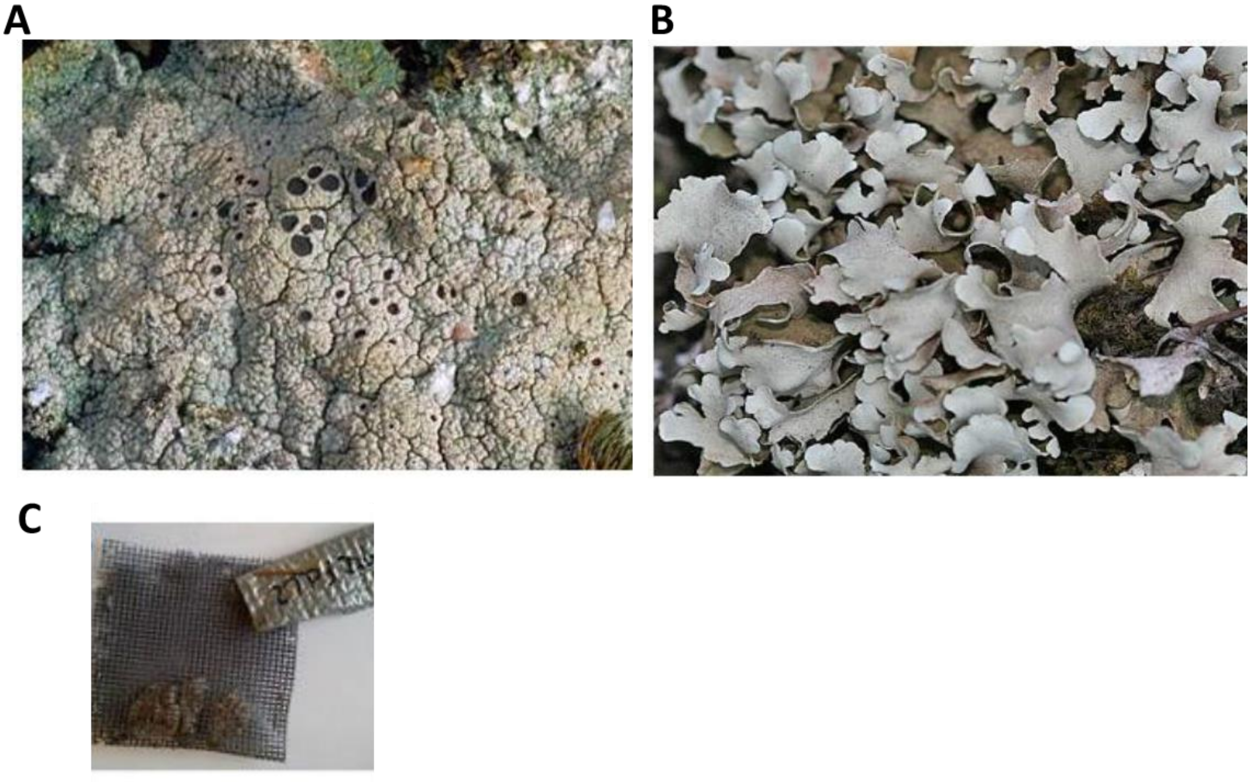

**Figure S2.** Photograph of the lichens used in the experiment. A) *Diploschistes diacapsis*; B) *Cladonia convoluta*; C) litterbags prepared for the experiment, right UV-block treatment; left UV-pass treatment. Source: Leif & Anita Stridval (http://www.stridvall.se) (B) y Stephen Sharnoff (http://www.sharnoffphotos.com) (A).

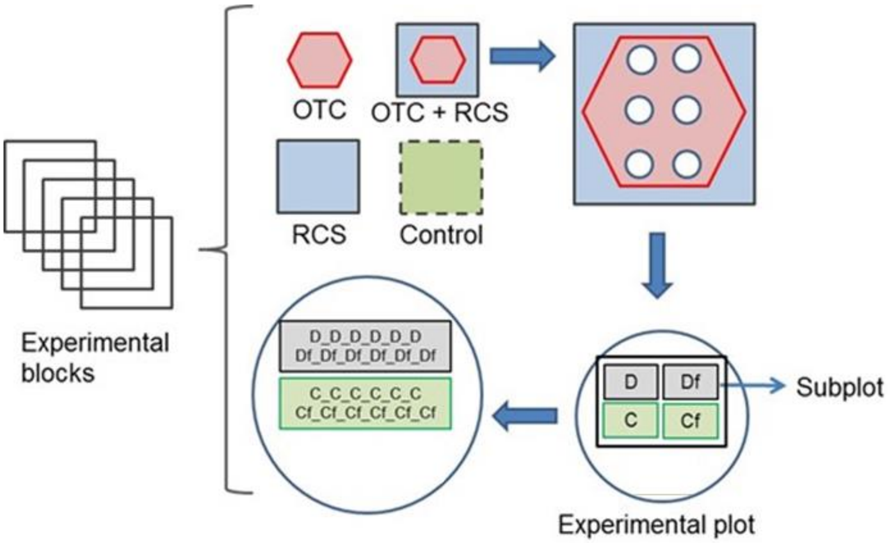


**Figure S3.** Experimental block scheme (n = 5) with three climate manipulation treatments (W, RE, REW) and control (C) with UV radiation treatment (UV-block or UV-pass) and litter lichen type (*D. diacapsis* or *C. convoluta*) and mixture of both lichen species (Mix).

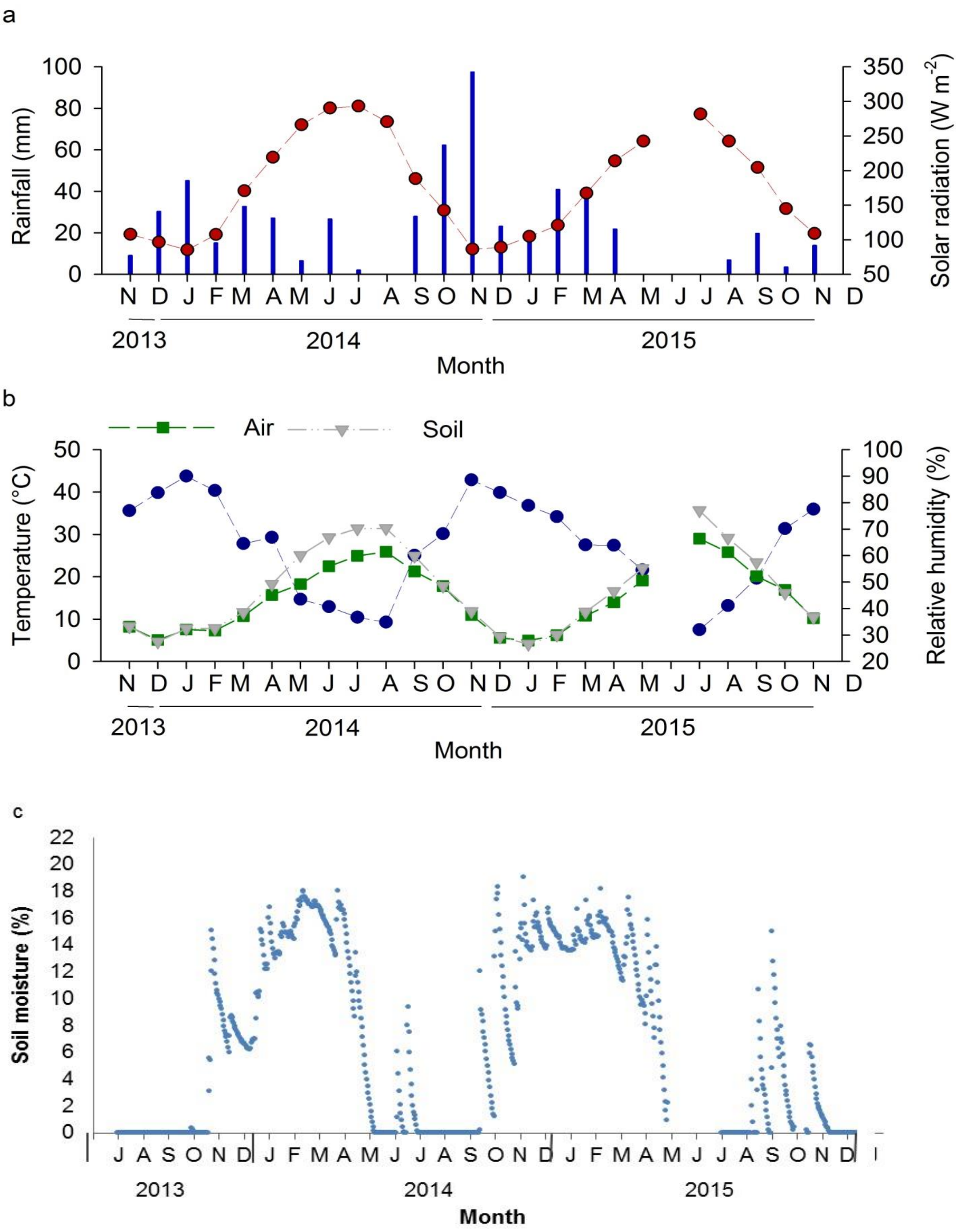


**Figure S4**. Monthly average of rain and solar radiation (a), air and soil temperature and air relative humidity (b) and soil moisture (c) at the study site throughout the course of the experiment.

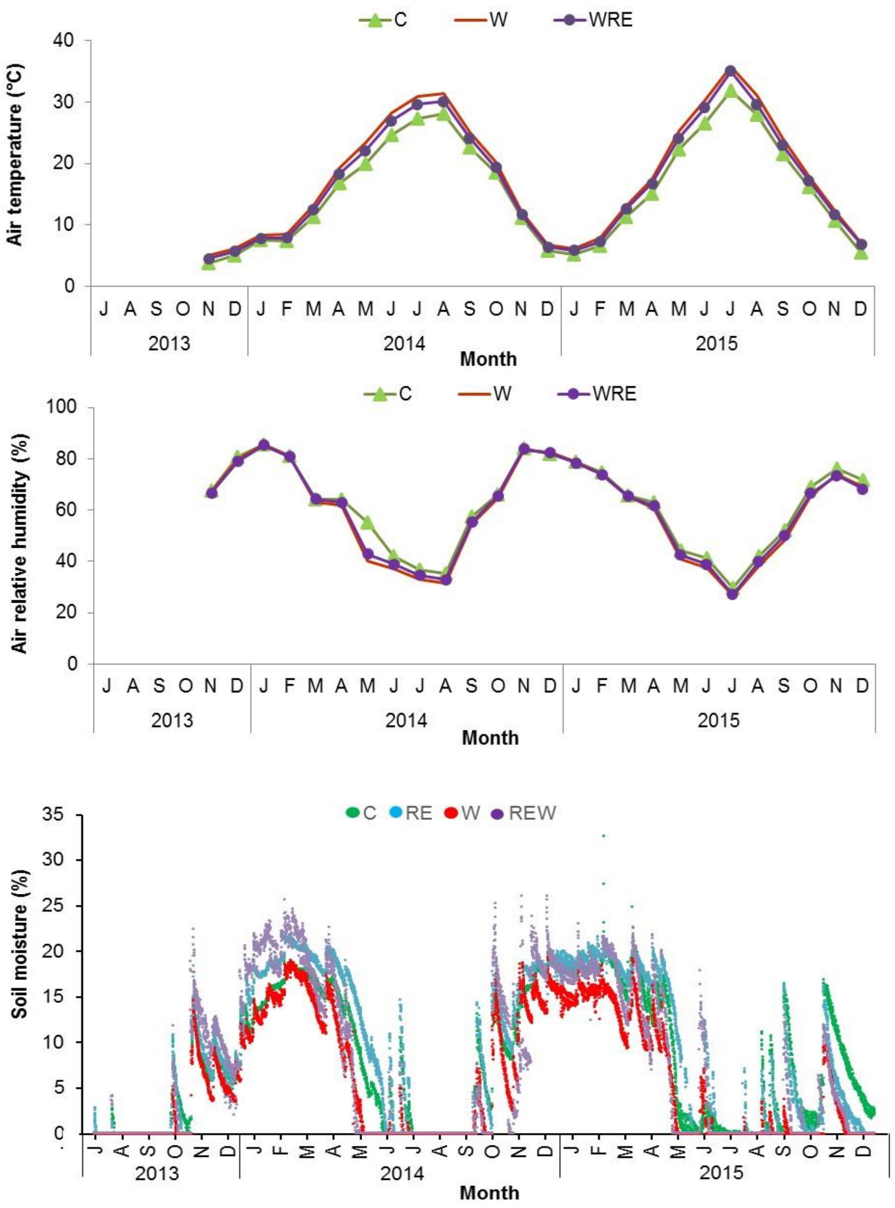


ç

**Figure S5.** Air temperature, air relative humidity (means of four sensors) and, soil moisture (mean of two sensors) over the duration of the study period (July 2013-December 2015) in the different climatic treatments: control (C), warming (W), rainfall exclusion (RE) and warming and rainfall exclusion (WRE) from a continuous experimental area. Standard errors were omitted for clarity.

**Table S1**. Decomposition litter decay (*k*), dry mass remaining (MR) chemical content (TOC: total organic content, N content and C/N ratio) of two lichen litter: *D. diacapsis* and *C. convoluta* near a year (intermediate) and two and a half year (end) of decomposition process started. Mean and SE are showed (n=5).

| | | | | | | Time | | | | | | | | | | | | | | | |
|---|---|---|---|---|---|---|---|---|---|---|---|---|---|---|---|---|---|---|---|---|---|
| | | | | | | Intermediate | | | | | | | | End | | | | | | | |
| | | | | $k$ (day$^{-1}$) | | MR (mg g-1) | | TOC (mg g-1) | | N (mg g-1) | | C/N | | MR (mg g-1) | | TOC (mg g-1) | | N (mg g-1) | | C/N | |
| Species | Main Effects and Interaction | | | Mean | SE | Mean | SE | Mean | SE | Mean | SE | Mean | SE | Mean | SE | Mean | SE | Mean | SE | Mean | SE |
| *D. diacapsis* | Treatclimate | C | | 0.0009 ± | 0.0002 | 85.14 ± | 1.69 | 129.11 ± | 13.43 | 5.11 ± | 0.31 | 25.96 ± | 3.50 | 54.07 ± | 7.69 | 139.52 ± | 3.13 | 6.04 ± | 0.31 | 23.29 ± | 0.73 |
| | | RE | | 0.0006 ± | 0.0001 | 81.36 ± | 4.10 | 109.57 ± | 5.60 | 6.82 ± | 0.31 | 16.08 ± | 0.49 | 60.62 ± | 6.63 | 121.13 ± | 10.23 | 7.45 ± | 0.47 | 16.70 ± | 1.91 |
| | | W | | 0.0013 ± | 0.0003 | 81.28 ± | 2.59 | 99.38 ± | 3.92 | 4.78 ± | 0.32 | 21.37 ± | 1.96 | 32.71 ± | 7.58 | 113.99 ± | 9.05 | 4.48 ± | 0.41 | 26.13 ± | 2.52 |
| | | WRE | | 0.0010 ± | 0.0002 | 82.26 ± | 2.47 | 101.69 ± | 8.79 | 4.59 ± | 0.31 | 22.82 ± | 2.83 | 37.20 ± | 5.46 | 97.50 ± | 5.28 | 4.14 ± | 0.27 | 23.73 ± | 0.93 |
| | Treatlight | UV-block | | 0.0008 ± | 0.0001 | 81.96 ± | 1.95 | 117.98 ± | 7.39 | 5.21 ± | 0.36 | 23.89 ± | 2.26 | 47.09 ± | 3.94 | 112.59 ± | 6.25 | 5.78 ± | 0.45 | 20.43 ± | 1.33 |
| | | UV-pass | | 0.0011 ± | 0.0002 | 82.74 ± | 2.21 | 101.89 ± | 5.27 | 5.44 ± | 0.32 | 19.22 ± | 1.32 | 46.47 ± | 7.08 | 123.48 ± | 6.92 | 5.28 ± | 0.48 | 24.50 ± | 1.51 |
| | Treatlight*Treatclimate | UV-block | C | 0.0011 ± | 0.0004 | 84.68 ± | 2.99 | 156.73 ± | 7.78 | 4.84 ± | 0.58 | 33.08 ± | 2.92 | 46.59 ± | 10.28 | 145.32 ± | 2.10 | 6.66 ± | 0.27 | 21.86 ± | 0.63 |
| | | UV-block | RE | 0.0007 ± | 0.0001 | 80.68 ± | 3.85 | 102.86 ± | 6.62 | 6.79 ± | 0.51 | 15.17 ± | 0.36 | 47.31 ± | 7.96 | 102.86 ± | 7.22 | 7.70 ± | 0.35 | 13.51 ± | 1.62 |
| | | UV-block | W | 0.0006 ± | 0.0001 | 79.50 ± | 4.91 | 99.12 ± | 5.55 | 4.58 ± | 0.30 | 21.76 ± | 1.39 | 46.13 ± | 7.83 | 96.33 ± | 4.67 | 4.20 ± | 0.06 | 22.98 ± | 1.42 |
| | | UV-block | WRE | 0.0007 ± | 0.0001 | 83.92 ± | 4.06 | 113.23 ± | 3.28 | 4.65 ± | 0.65 | 25.54 ± | 4.26 | 48.98 ± | 1.65 | 105.87 ± | 6.56 | 4.54 ± | 0.28 | 23.36 ± | 1.14 |
| | | UV-pass | C | 0.0007 ± | 0.0003 | 85.74 ± | 1.37 | 101.49 ± | 8.89 | 5.39 ± | 0.24 | 18.84 ± | 1.46 | 61.55 ± | 11.52 | 133.72 ± | 3.32 | 5.41 ± | 0.15 | 24.73 ± | 0.48 |
| | | UV-pass | RE | 0.0005 ± | 0.0002 | 82.21 ± | 8.69 | 116.28 ± | 8.23 | 6.85 ± | 0.48 | 16.99 ± | 0.51 | 70.60 ± | 6.68 | 139.41 ± | 11.71 | 7.20 ± | 0.95 | 19.88 ± | 2.35 |
| | | UV-pass | W | 0.0020 ± | 0.0005 | 83.05 ± | 2.10 | 99.64 ± | 6.77 | 4.98 ± | 0.60 | 20.98 ± | 4.14 | 19.29 ± | 9.32 | 131.65 ± | 8.69 | 4.76 ± | 0.87 | 29.27 ± | 4.44 |
| | | UV-pass | WRE | 0.0012 ± | 0.0003 | 80.60 ± | 3.20 | 90.15 ± | 15.58 | 4.53 ± | 0.24 | 20.09 ± | 3.80 | 25.42 ± | 2.77 | 89.12 ± | 5.12 | 3.74 ± | 0.36 | 24.11 ± | 1.68 |
| *C. convoluta* | Treatclimate | C | | 0.0005 ± | 0.0001 | 82.15 ± | 3.19 | 374.33 ± | 10.87 | 5.02 ± | 0.25 | 75.96 ± | 5.75 | 60.43 ± | 5.67 | 372.10 ± | 11.87 | 6.07 ± | 0.38 | 62.42 ± | 4.16 |
| | | RE | | 0.0005 ± | 0.0001 | 83.76 ± | 2.20 | 380.69 ± | 15.30 | 4.86 ± | 0.34 | 79.75 ± | 4.64 | 61.44 ± | 5.50 | 328.60 ± | 7.52 | 5.89 ± | 0.41 | 57.04 ± | 3.75 |
| | | W | | 0.0006 ± | 0.0001 | 78.77 ± | 2.58 | 355.44 ± | 10.94 | 5.51 ± | 0.39 | 66.52 ± | 6.00 | 58.19 ± | 5.01 | 328.76 ± | 11.24 | 5.15 ± | 0.23 | 64.22 ± | 2.33 |
| | | WRE | | 0.0008 ± | 0.0001 | 76.58 ± | 2.92 | 367.33 ± | 24.53 | 5.44 ± | 0.27 | 68.45 ± | 5.51 | 48.23 ± | 5.29 | 346.95 ± | 10.61 | 5.40 ± | 0.33 | 65.82 ± | 5.48 |
| | Treatlight | UV-block | | 0.0007 ± | 0.0001 | 78.34 ± | 1.92 | 368.02 ± | 10.93 | 5.08 ± | 0.19 | 73.95 ± | 4.05 | 49.46 ± | 3.38 | 336.54 ± | 8.39 | 5.95 ± | 0.29 | 57.95 ± | 2.83 |
| | | UV-pass | | 0.0005 ± | 0.0001 | 82.94 ± | 1.88 | 370.87 ± | 11.84 | 5.34 ± | 0.26 | 71.39 ± | 4.01 | 64.46 ± | 3.55 | 351.66 ± | 8.72 | 5.31 ± | 0.16 | 66.80 ± | 2.36 |
| | Treatlight*Treatclimate | UV-block | C | 0.0006 ± | 0.0001 | 78.03 ± | 4.28 | 393.93 ± | 5.15 | 4.51 ± | 0.15 | 87.52 ± | 3.52 | 52.94 ± | 7.16 | 376.62 ± | 9.00 | 6.38 ± | 0.66 | 60.70 ± | 7.92 |
| | | UV-block | RE | 0.0006 ± | 0.0002 | 84.28 ± | 2.54 | 399.40 ± | 5.84 | 5.09 ± | 0.28 | 78.78 ± | 3.27 | 51.28 ± | 9.52 | 329.61 ± | 5.61 | 6.67 ± | 0.45 | 49.94 ± | 4.02 |
| | | UV-block | W | 0.0007 ± | 0.0001 | 77.31 ± | 3.80 | 365.03 ± | 14.67 | 5.20 ± | 0.35 | 71.35 ± | 8.15 | 49.94 ± | 4.71 | 311.29 ± | 15.71 | 4.69 ± | 0.14 | 66.46 ± | 3.24 |
| | | UV-block | WRE | 0.0009 ± | 0.0001 | 72.57 ± | 3.60 | 313.70 ± | 6.70 | 5.51 ± | 0.55 | 58.17 ± | 6.29 | 42.69 ± | 7.59 | 328.64 ± | 3.55 | 6.05 ± | 0.31 | 54.68 ± | 3.14 |
| | | UV-pass | C | 0.0004 ± | 0.0001 | 87.30 ± | 3.83 | 354.73 ± | 13.40 | 5.53 ± | 0.19 | 64.40 ± | 4.43 | 67.91 ± | 8.09 | 367.57 ± | 24.57 | 5.76 ± | 0.43 | 64.15 ± | 4.57 |
| | | UV-pass | RE | 0.0004 ± | 0.0001 | 83.24 ± | 3.90 | 361.97 ± | 28.05 | 4.63 ± | 0.67 | 80.73 ± | 9.80 | 69.57 ± | 4.18 | 327.60 ± | 15.81 | 5.11 ± | 0.18 | 64.13 ± | 1.90 |
| | | UV-pass | W | 0.0004 ± | 0.0001 | 80.58 ± | 3.73 | 345.84 ± | 17.05 | 5.82 ± | 0.75 | 61.70 ± | 9.50 | 68.51 ± | 7.04 | 346.23 ± | 8.92 | 5.61 ± | 0.18 | 61.97 ± | 3.41 |
| | | UV-pass | WRE | 0.0007 ± | 0.0001 | 80.58 ± | 4.01 | 420.95 ± | 9.42 | 5.36 ± | 0.20 | 78.73 ± | 2.55 | 52.65 ± | 7.44 | 365.25 ± | 14.65 | 4.76 ± | 0.17 | 76.96 ± | 4.05 |

**Table S2.** Summary statistics (three-way ANOVA) of treatment effects and their interactions on mass remaining (%) at the 12 and 30 months of decomposition experiment, and decay constant (day$^{-1}$) is also display. Bold numbers kept statistical significant (P < 0.05).

| | | | Time | | | |
|---|---|---|---|---|---|---|
| | | | Intermediate | | Final | |
| **Variable** | **Effect** | **df** | ***F*** | ***P*** | ***F*** | ***P*** |
| Mass Remaining | Species | 1,52 | 0.99 | 0.325 | 7.58 | **0.008** |
| | Light | 1,52 | 1.81 | 0.185 | 2.33 | 0.133 |
| | Climate | 3,52 | 0.99 | 0.403 | 4.16 | **0.010** |
| | Species * Light | 1,52 | 1.01 | 0.320 | 5.16 | **0.027** |
| | Species * Climate | 3,52 | 0.65 | 0.588 | 1.81 | 0.157 |
| | Light * Climate | 3,52 | 0.25 | 0.863 | 2.77 | 0.051 |
| | Species * Light * Climate | 3,52 | 0.64 | 0.590 | 2.36 | 0.082 |
| Decay constant | Species | 1,64 | | | 9.20 | **0.018** |
| | Light | 1,64 | | | 0.19 | 0.191 |
| | Climate | 3,64 | | | 2.46 | **0.038** |
| | Species * Light | 1,64 | | | 6.84 | **0.014** |
| | Species * Climate | 3,64 | | | 1.46 | 0.516 |
| | Light * Climate | 3,64 | | | 3.25 | 0.093 |
| | Species * Light * Climate | 3,64 | | | 4.21 | **0.026** |

**Table S3**. Pairwise comparisons by LSD test of Species * Light and Climate * Light interactions effects in % Mass Remaining variable at the end of the experiment (month 30).

| Effect | Factors | | | Mean difference | *P* |
|---|---|---|---|---|---|
| Species * Light | UV-block | *D. diacapsis* vs. | *C. convoluta* | -1.964 | .739 |
| | UV-pass | | | -20.442 | **.001** |
| | Diploschistes | UV-block vs. | UV-pass | 3.035 | .618 |
| | Cladonia | | | -15.444 | **.007** |
| Climate * Light | C | UV-block vs. | UV-pass | -14.962 | **.048** |
| | W | | | 4.136 | .609 |
| | RE | | | -20.787 | **.016** |
| | WRE | | | 6.794 | .439 |
| | UV-block | C vs. | W | 1.730 | .821 |
| | | | RE | .469 | .954 |
| | | | WRE | 3.932 | .632 |
| | | W vs. | RE | -1.262 | .881 |
| | | | WRE | 2.201 | .794 |
| | | RE vs. | WRE | 3.463 | .699 |
| | UV-pass | C vs. | W | 20.828 | **.010** |
| | | | RE | -5.356 | .484 |
| | | | WRE | 25.687 | **.002** |
| | | W vs. | RE | -26.185 | **.002** |
| | | | WRE | 4.859 | .564 |
| | | RE vs. | WRE | 31.044 | **.000** |

**Table S4**. Total organic carbon (TOC) and total nitrogen (TN ) concentrations (mg $g^{-1}$) of litter of both species *Diploschistes diacapsis* and *Cladonia convoluta* at the beginning of the experiment. Different letters show significant differences between species within each row after a Student t test ($P < 0.05$). Data are means ± SE (n = 6).

| | *D. diacapsis* | | | *C. convoluta* | | |
|---|---|---|---|---|---|---|
| | Mean | | SE | Mean | | SE |
| OC | 138.9 | ± | 10.0a | 371.7 | ± | 9.50b |
| TN | 4.6 | ± | 0.29a | 6.5 | ± | 0.45b |
| C/N | 30.0 | ± | 1.62a | 58.6 | ± | 4.63b |

**Table S5.** Three-way ANOVA results of main effects and their interactions on chemical composition of the lichen-litter at xx and 30 months after initialization of the decomposition experiment. *P* values below 0.05 are in bold.

| | | | Time | | | |
|---|---|---|---|---|---|---|
| | | | Intermediate | | Final | |
| **Variable** | **Effect** | **df** | ***F*** | ***P*** | ***F*** | ***P*** |
| OC | Species | 1 | 1025.78 | <**0.001** | 4324.15 | < **0.001** |
| | Light | 1 | 0.67 | 0.420 | 14.30 | < **0.001** |
| | Climate | 3 | 1.79 | 0.168 | 23.20 | < **0.001** |
| | Species * Light | 1 | 1.37 | 0.250 | 0.38 | 0.539 |
| | Species * Climate | 3 | 0.49 | 0.690 | 7.48 | < **0.001** |
| | Light * Climate | 3 | 5.15 | **0.005** | 7.49 | < **0.001** |
| | Species * Light * Climate | 3 | 11.14 | < **0.001** | 7.52 | < **0.001** |
| TN | Species | 1 | 0.28 | 0.600 | 0.53 | 0.469 |
| | Light | 1 | 1.14 | 0.292 | 16.57 | < **0.001** |
| | Climate | 3 | 2.99 | **0.044** | 45.50 | < **0.001** |
| | Species * UV | 1 | 0.01 | 0.941 | 0.24 | 0.626 |
| | Species * Climate | 3 | 8.39 | <**0.001** | 9.76 | < **0.001** |
| | Light * Climate | 3 | 1.19 | 0.330 | 18.96 | < **0.001** |
| | Species * Light * Climate | 3 | 0.21 | 0.890 | 1.92 | 0.125 |
| C/N | Species | 1 | 361.41 | < **0.001** | 546.64 | < **0.001** |
| | Light | 1 | 1.81 | 0.187 | 14.34 | **0.001** |
| | Climate | 3 | 1.28 | 0.297 | 5.05 | **0.006** |
| | Species * Light | 1 | 0.15 | 0.699 | 1.96 | 0.171 |
| | Species * Climate | 3 | 2.59 | 0.068 | 0.25 | 0.859 |
| | Light * Climate | 3 | 4.43 | **0.010** | 2.34 | 0.091 |
| | Species * Light * Climate | 3 | 2.69 | 0.063 | 3.93 | **0.017** |

**Table S7**. Two-way ANOVA at the end of decomposition process of two litter lichen species (expressed as species) for the four climate change treatments but only for the UV-block conditions. $P < 0.05$ are in bold.

| Variable | Factor | df | F | *P* |
|---|---|---|---|---|
| Mass remaining | Species | 2,40 | 0.07 | 0.997 |
| | Climate | 3,40 | 0.23 | 0.877 |
| | Species * Climate | 6,40 | 0.18 | 0.981 |
| Decomposition rate ($day^{-1}$) | Species | 2,48 | 0.73 | 0.723 |
| | Climate | 3,48 | 1.14 | 0.296 |
| | Species * Climate | 6,48 | 1.21 | 0.512 |
| TOC ($mg\ g^{-1}$) | Species | 2,24 | 758.32 | **< 0.001** |
| | Climate | 3,24 | 8.48 | **0.001** |
| | Species * Climate | 6,24 | 9.71 | **< 0.001** |
| TNC ($mg\ g^{-1}$) | Species | 2,24 | 3.38 | 0.051 |
| | Climate | 3,24 | 23.24 | **< 0.001** |
| | Species * Climate | 6,24 | 1.86 | 0.129 |
| C:N | Species | 2,24 | 64.23 | **< 0.001** |
| | Climate | 3,24 | 4.34 | **0.014** |
| | Species * Climate | 6,24 | 1.13 | 0.377 |
